# The Profiling of Bisecting N-acetylglucosamine (GlcNAc) Modification in Human Amniotic Membrane by Glycomic and Glycoproteomic Analyses

**DOI:** 10.1101/2020.06.09.141168

**Authors:** Qiushi Chen, Yuanliang Zhang, Keren Zhang, Jie Liu, Huozhen Pan, Xinran Wang, Siqi Li, Dandan Hu, Zhilong Lin, Yun Zhao, Guixue Hou, Feng Guan, Hong Li, Siqi Liu, Yan Ren

**Author notes:** Equal contribution (Qiushi Chen, Yuanliang Zhang, Keren Zhang). Corresponding authors (Liu S), (Ren Y).

## Abstract

It is acknowledged that the bisecting N-acetylglucosamine (GlcNAc) structure, a GlcNAc linked to the core β-mannose residue via a β1,4 linkage, represents a special type of N-glycosylated modification and has been reported to be involved in various biological processes, such as cell adhesion and fetal development. Clark et al. has found that the majority of N-glycans in human trophoblasts bearing a bisecting GlcNAc. This type of glycan has been reported to help trophoblasts get resistant to natural killer (NK) cell-mediated cytotoxicity, and this would provide a possible explanation for the question how could the mother nourish a fetus within herself without rejection. Herein, we hypothesized that human amniotic membrane which is the last barrier for the fetus may also express bisecting type glycans to protect the fetus. To test this hypothesis, glycomic analysis of human amniotic membrane was performed, and the bisecting N-glycans with high abundance were detected. In addition, we re-analyzed our proteomic data with high fractionation and amino acid sequence coverage from human amniotic membrane, which had been released for the exploration of human missing proteins. The presence of bisecting GlcNAc peptides was revealed and confirmed. A total of 41 glycoproteins with 43 glycopeptides were found to possess a bisecting GlcNAc, 25 of which are for the first time to be reported to have this type of modification. These results provide the profiling of bisecting GlcNAc modification in human amniotic membrane and benefit to the function studies of glycoproteins with bisecting GlcNAc modification and the function studies in immune suppression of human placenta. The mass spectrometry placenta data are available via ProteomeXchange (PXD010630).

## Introduction

A distinctive structural feature of N-glycans is the presence of several N-acetylglucosamine (GlcNAc) antennae that are sequentially synthesized by a group of Golgi-resident glycosyltransferases, N-acetylglucosaminyltransferases (GlcNAc-Ts) [1, 2]. There are three categories of N-glycans: high-mannose, hybrid and complex. Hybrid and complex N-glycans may carry a bisecting GlcNAc group, which forms a new subtype of glycan termed bisected glycan [3, 4]. This type of glycan was reported in the 1970s and was detected by a combination of sequential exoglycosidase digestion, methylation derivatization, acetolysis and Smith degradation from ovalbumin [5]. GlcNAc was transferred to the 4-position of the β-linked core mannose (Man) residue in complex or hybrid N-glycans by the β1,4-mannosyl-glycoprotein 4-β-N-acetylglucosaminyltransferase (GlcNAc-T III), however, this GlcNAc is usually not considered as an antenna initiation point as it cannot be further extended [1, 3, 6-8].

GlcNAc-T III is encoded by the gene *mgat3*, which was initially discovered from hen oviducts in 1982 [8]. The existence of a bisecting GlcNAc prevents α-mannosidase II from trimming and has been proved to inhibit the activities of GlcNAc-T II, GlcNAc-T IV and GlcNAc-T V *in vitro* as well [1, 3, 9]. The addition of bisecting GlcNAc confers unique lectin recognition properties to this subtype of glycan [7, 10]. The B16 mouse melanoma transfected by *mgat3* shows weaker binding to phytohemagglutinin-L (PHA-L) but stronger binding to Phaseolus vulgaris erythroagglutinin (PHA-E). The lectins of PHA-L and PHA-E show specific recognition to multiple antennary glycans and bisecting GlcNAc structures, respectively [3, 11].

The bisecting GlcNAc is essential for many biological processes, including tumor development and immune response [10, 12]. For instance, bisecting GlcNAc structures have been reported to possess immune suppression functions. Human K562 cells are easily killed by natural killer (NK) cells; however, after being transfected with the gene *mgat3*, K562 cells become NK cell resistant due to the expression of bisecting GlcNAc [13-15].

In the 1990s Clark et al. proposed the human fetoembryonic defense system (hu-FEDS) hypothesis to answer the question how the mother nourishes a fetus within herself for several months without rejection. This question was initially raised by Sir Peter Brian Medawar who shared 1960 Nobel Prize in Physiology or Medicine with Sir Frank Macfarlane Burnet [16, 17]. Since the hypothesis was conceived, it has been intensively tested. An increasing amount of evidence has been shown to support this hypothesis [18, 19]. In 2016, Clark et al. has found that the functional glycan structure (bisecting GlcNAc) that are present on human gametes are also expressed on human trophoblasts, and more importantly, the majority of N-glycans in human trophoblasts possesses a bisecting GlcNAc [6, 20]. It is also reported in 2016 that the occurrence of N-glycans with a bisecting GlcNAc modification on glycoproteins has many implications in immune biology [21].

The traditional method that used for bisecting GlcNAc detection is based on the lectin recognition of *Phaseolus vulgaris* erythroagglutinin (PHA-E); however, the poor binding specificity hinders the application of this method [22]. With the development of mass spectrometry (MS) with high resolution plus improved sensitivity and accuracy, MS-based analysis has provided precise characterization for glycosylations [6, 23-25].

Various MS-based approaches have been released and proved as efficient tools for bisecting GlcNAc modification studies [6, 20, 22, 23, 26]. These bisecting GlcNAc determination approaches can be divided into two catogories based on the detection targets, namely, glycan and glycopeptide. The basic idea of the glycan method is to determine the presence of 3,4,6-linked Man, which can be achived via either MS7 analysis [23] or partially methylated alditol acetates (PMAA) derivatization together with gas chromatography-mass spectrometry (GC-MS) [6, 20, 26]. The fundamental idea of the glycopeptide method is to determine the presence of the characteristic fragment ion(s) [Peptide+HexNAc_3_Hex] or [Peptide+FucHexNAc_3_Hex] or both, and this can be achived through MS/MS fragmentation under low energy collisions, which has been described in a recently published paper in Analytical Chemistry in 2019 [22].

Amniotic membrane (amnion) is the innermost layer of placenta. It is the last barrier for the fetus. In addition, amnion membrane has been considered to be a potential stem cell reservoir with wide applications in periodontics, tissue regeneration and surgery [27-29]. Knowing more about amniotic membrane is getting increasingly worthwhile. As a matter of fact, amniotic membrane is an optimal sample for analysis as the possibility of it being contaminated by blood cells, neurocytes and lymphocytes are low if the membrane is washed in saline or PBS prior to enzyme digestion [30]. More importantly, by far the glycomic and glycoproteomic studies on amnion have not been reported yet. Herein, we hypothesized that human amniotic membrane may also express bisecting type glycans to protect the fetus. To test the hypothesis, we employed the MS-based approach to determine the presence of bisecting GlcNAc on amniotic membrane proteins. In addition, the glycosylation sites of the corresponding glycoproteins were determined.

## Results and Discussion

### Preparation of human amniotic membrane and determination of protein concentration

Human amniotic membrane was isolated from a placenta (Figure 1A-C) accordingly [30]. The proteins were extracted by 8M urea as reported before [31]. SDS-PAGE image was used to check the extraction efficiency and whether plasma protein contamination existed (Figure 1D).

**Figure 1.**
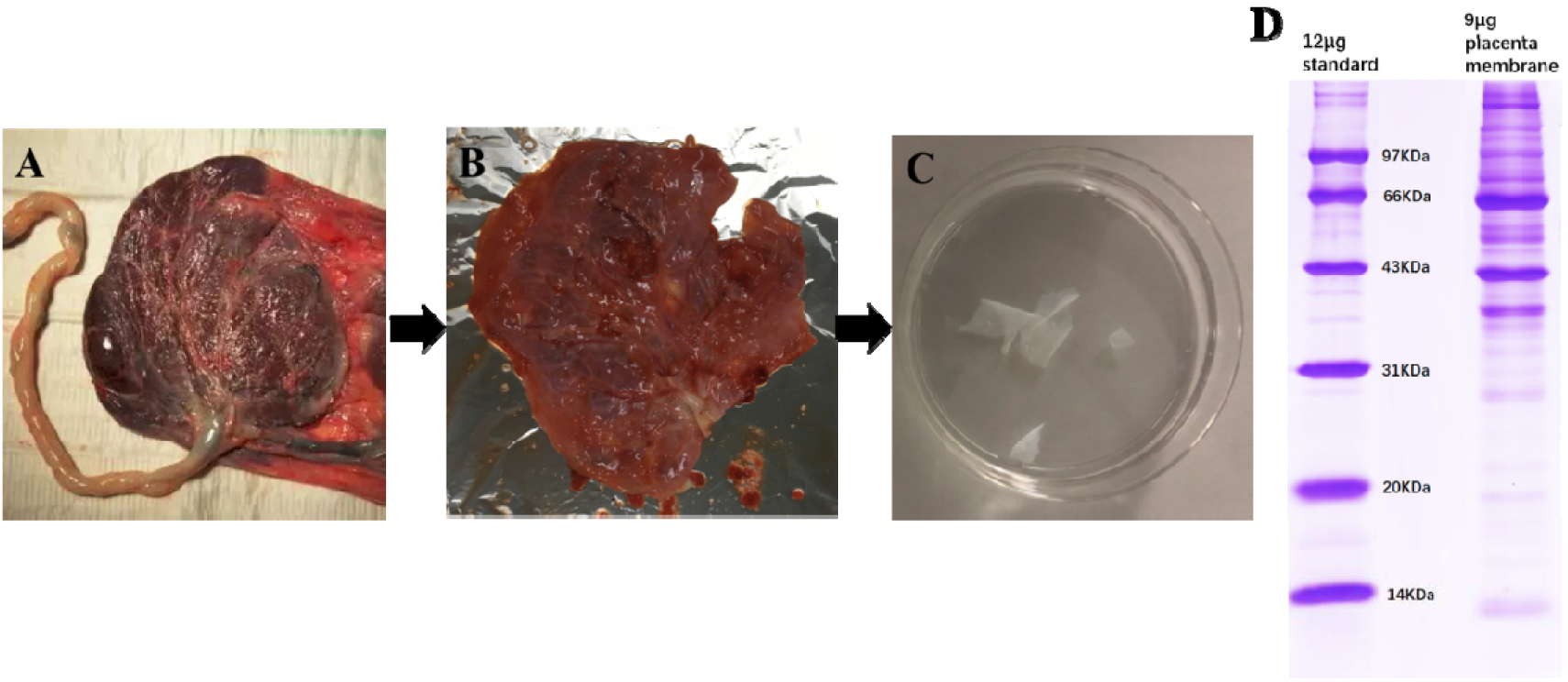
The preparation and protein extraction of human amniotic membrane A. human placenta and umbilical cord; B. chorion and amniotic membrane section; C. amniotic membrane after washing with PBS; D. SDS-PAGE image for the extracted proteins from human amniotic membrane

### MALDI-TOF MS analysis of the N-glycans from human amniotic membrane

The N-linked glycans were released from the amniotic membrane proteins and permethylated for sensitive MALDI-TOF analysis, which were subjected to glycomic profiling analysis exactly as described previously [26, 32]. High quality MALDI-TOF data were obtained for these N-glycans. As shown in Figure 2, 45 N-glycans were detected; high mannose (e.g. m/z 1579.4 and m/z 1783.4) and complex glycans (e.g. m/z 2489.4 and m/z 2938.7) are present in the human amniotic membrane. several common features of mammalian cell N-glycomes [25, 33] were observed, such as core fucosylated GlcNAc (e.g. m/z 2693.7), N-acetylneuraminic acid (NeuAc) capped antennae (e.g. m/z 2605.6) and (N-acetyllactosamine) LacNAc units (e.g. m/z 3591.2) which in some cases form tandem repeats to yield oligo-LacNAc antennae. The amount of the complex glycans accounts for nearly 74% of all detected glycans. Approximately 80% of the complex-type glycans carry core α1–6 linked fucose, which is consistent with their localization to the plasma membrane. The m/z value of the most complex N-glycan observed is m/z 4279.9, which contains 19 monosaccharides. Importantly, The peak with m/z 2489.4 was found with the most abundance, which is speculated as a potential core-fucosylated biantennary bisecting modification. Some minor signals for N-glycans bearing potential Lewis type antenna, such as m/z 2418.3 and m/z 2837.7, were also perceived.

**Figure 2.**
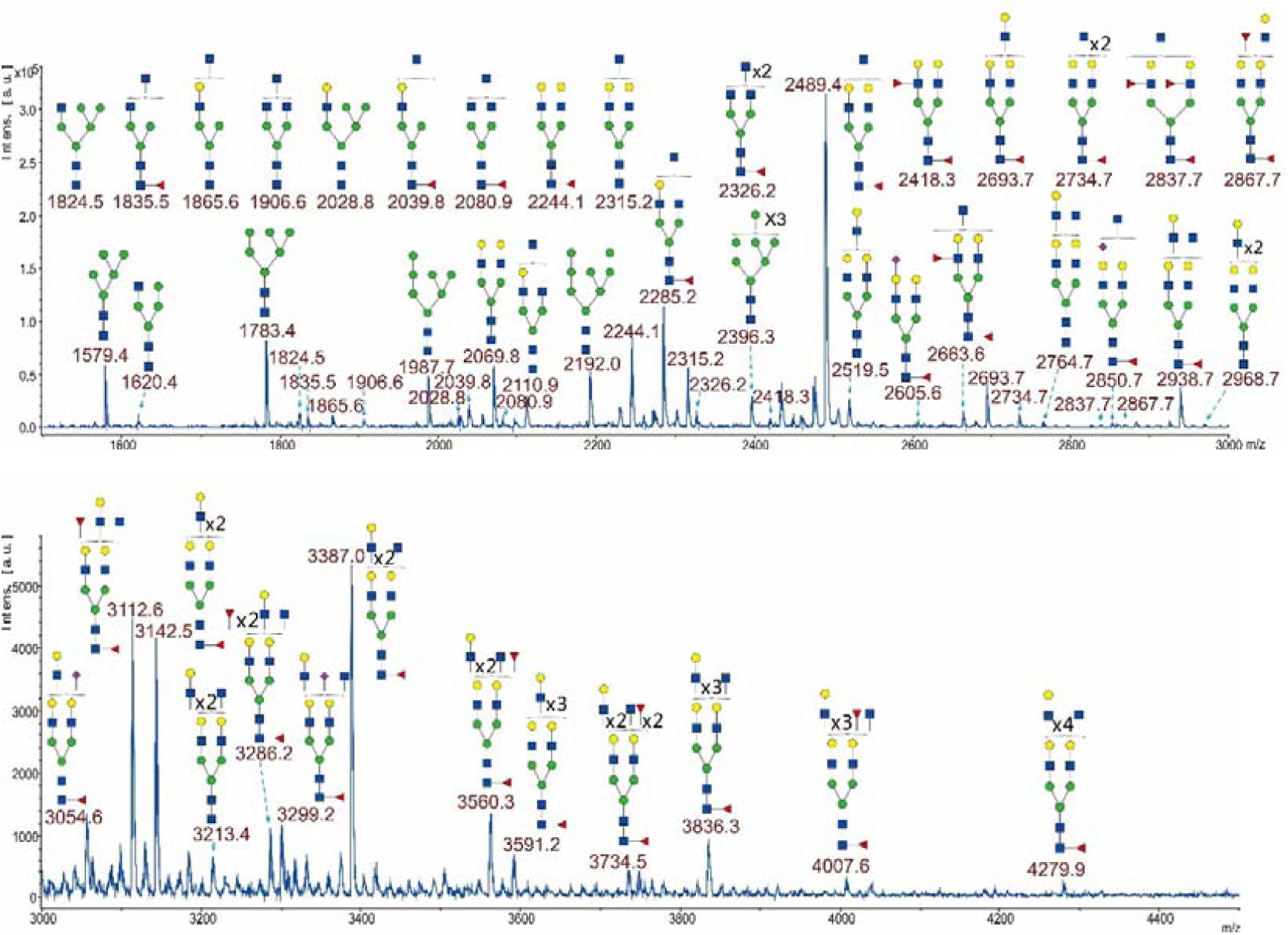
Annotated MALDI-TOF MS spectra of permethylated N-glycans from human amniotic membrane. The top panel shows the glycans in the mass range from m/z 1500 to 3000 and the bottom panel shows the glycans in the mass range from m/z 3000 to 4500. All ions are [M+Na]^+^. Peaks are labeled with their m/z values, Putative structures are described basing on the molecular weight and N-glycan biosynthetic pathway. Annotations are simplified to biantennary structures, with additional N-acetyllactosamine (LacNAc) units, Fucose (Fuc) and N-acetylglucosamine (GlcNAc) listed outside the bracket, ▪ GlcNAc, • Mannose (Man), • Galactose (Gal), ▴ Fuc, ♦ NeuAc.

### MS8 analysis of the N-glycan at m/z 2489.4 by an Orbitrap Fusion™ Lumos™ Tribrid™ mass spectrometer

The query glycans with higher abundance were further fragmentated under MSn (n=2-8) analysis mode to get their fine structures. The sequential collision of target glycans (m/z 2489.25) with the highest intensity and a potential bisected N-glycan structure were carried out to distinctively confirm the presence of bisecting GlcNAc. Figure 3 shows the logical order of the MS8 approach that could distinguish the presence of bisecting GlcNAc structure of the glycans from other glycan structures which possess the same m/z at 2489.25 in Obitrap lumos MS/MS analysis.

**Figure 3.**
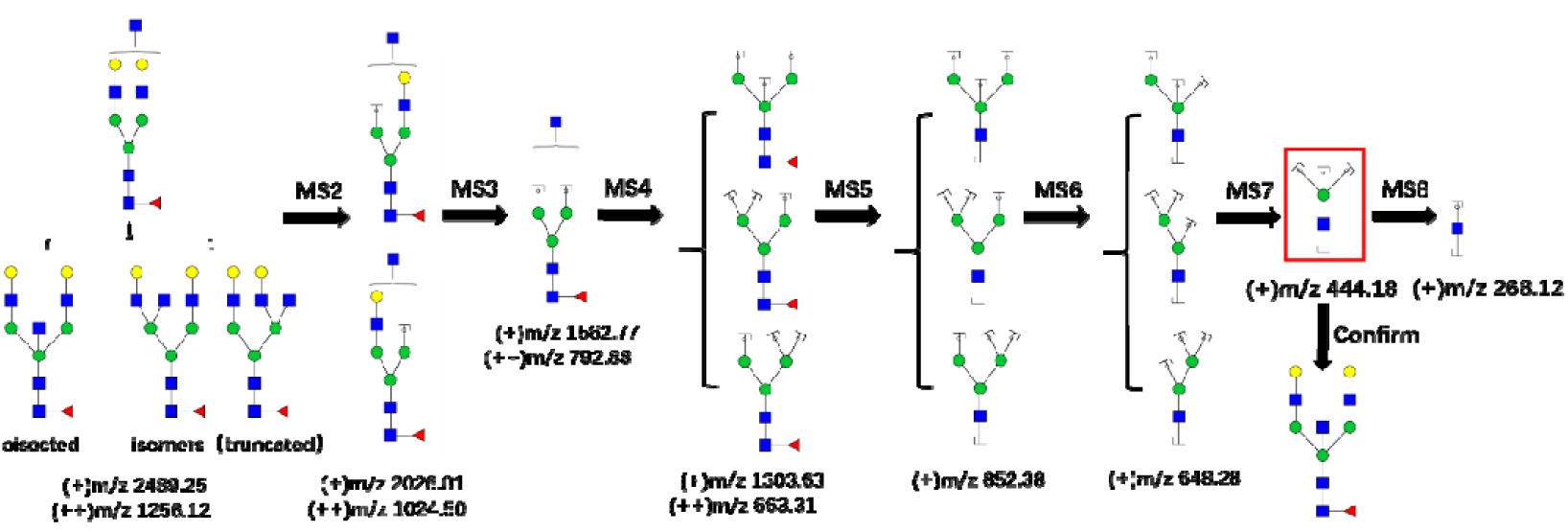
The MS8 approach to confirm the presence of bisected N-glycan. The fragment ion at m/z 444.18 (theoretical value) in the red frame is the characteristic ion of the bisecting type glycan. ▪ GlcNAc, • Man, • Gal, ▴Fuc.

Theoretically, the MS7 spectra of glycans at m/z 2489.25 should display the characteristic ions of bisecting GlcNAc glycans at m/z 444.18 and its MS8 spectra would be occupied by the peak of m/z 268.12 to support the presence of GlcNAc group. In fact, the MS7 scan (Figure 4) for the glycan fragment ions contains a dominant fragment ion at m/z 444.07, which is about 0.11 Dalton mass difference from its theoretical m/z value. Due to the low resolution of ion trap analyzer, the detected ion m/z values have a relatively high mass error, which is different from the MS1 and MS2 analysis under Fourier transform mass spectrometry (FTMS) analyzer [34, 35]. As shown in Figure 4, the characteristic fragment ion at m/z 444.07 is produced by m/z 647.93 (theoretical m/z: 648.28) via losing a Man BY ion (mannose, the green circle). In addition to this ion, three more ions were observed: m/z at 421.17 (same as the theoretical m/z value) is the resulting ion from m/z 444.07 losing a GlcNAc BZ ion, m/z at 403.13 (theoretical m/z :403.16) represents the resulting ion from m/z 444.07 losing a GlcNAc BY ion and m/z at 267.83 stands for a GlcNAc BY ion (theoretical m/z: 268.12). MS8 analysis was further performed to show that the ion at m/z 444.07 is a glycan fragment ion indeed, not an noise. All the other MSn (n=2-6, 8) spectra for the glycan can be found in Supplementary Figure 1.

**Figure 4.**
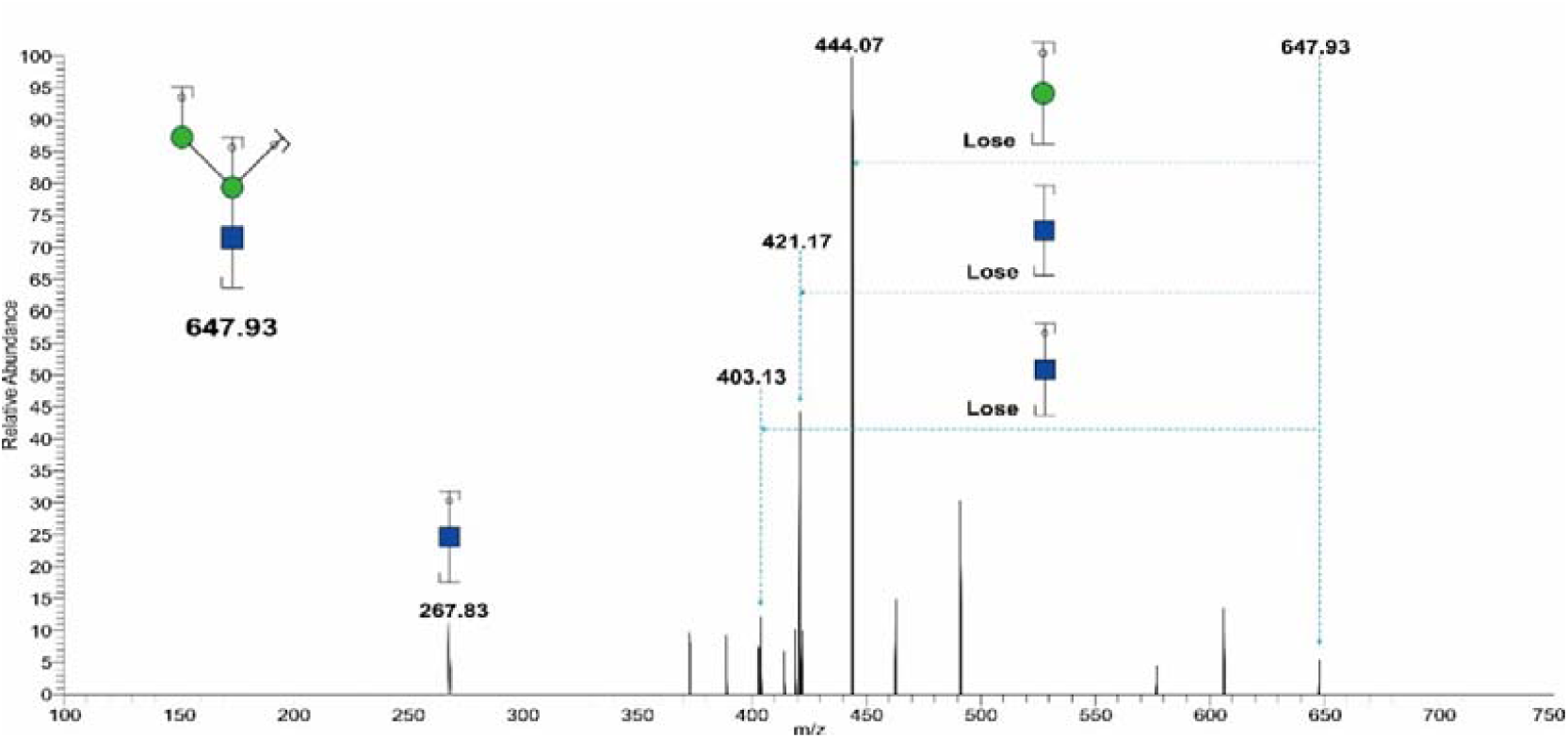
Annotated ESI ITMS MS7 spectrum of monocharged permethylated N-glycan fragment ion at m/z 647.93 from human amniotic membrane. Assignments of the possible fragment ions are indicated on the cartoons and on the spectrum. The number indicated above the peak is the m/z value of the fragment ion (resulting ion) that has been detected by the mass spectrometer. Data were acquired in the form of [M+Na]^+^. The energy under CID and HCD mode firstly breaks glycosidic bonds to form B ions or Y ions, further fragmentation will produce BY ions or YY ions or BB ions, and so on. ▪ GlcNAc, • Man.

Compared to other MS methods for detecting bisecting GlcNAc modification, our method does not require extra enzymatic treatments (e.g.β1,4-galactosyltransferase) or extra derivatization procedures (e.g. partially methylated alditol acetates derivatization) or lectin recognitions (e.g. phaseolus vulgaris erythroagglutinin), which makes this method a straightforward approach for bisecting GlcNAc determination [6, 10, 26, 36]. It could greatly facilitate the researches on bisected N-glycans.

### Glycoproteomic analysis for human amniotic membrane proteins

Since the presence of bisecting GlcNAc modification has been confirmed from the human amniotic membrane proteins, therefore it is necessary to find the origin or the location of this type of glycan modification in proteins. We have reported an extensive proteomic analysis to human amniotic membranes, in which the peptides were fractionated by a three-dimensional separation approach according to their hydrophobicity. Thus, the data for 40 peptide fractions with each injection of 2-hour MS run were collected. The deep analysis resulted in 9941 proteins with 163091 peptides detected from human amniotic membranes, which offered a good library with higher peptide sequence coverage for the glycopeptide searching. Searching glycopeptides from the library were mainly based on the combination of Proteome Discoverer and Xcalibur software to figure out the glycopeptides and manually checking of oxonium ions, monosaccharide and oligosaccharide neutral loss patterns in MS2 spectra to get the corresponding glycan structure.

With the confident analysis, 43 glycopeptides belongling to 41 glycoproteins were found to possess bisected N-glycans, in which the glycosylation sites and linked glycans were confirmed by manual checking with the presence of either peptide+HexNAc_3_Hex or peptide+FucHexNAc_3_Hex or both of them as solid evidences. Supplementary Table 1 summarizes all the glycoproteins with bisecting N-glycan modification, including their accession numbers, protein names, peptides modified by bisecting GlcNAc glycans and cellular locations. All of the glycoproteins were assigned as membrane proteins or extracellular matrix proteins by searching them in UniProt library, in which 18 and 22 proteins belong to cell membrane and extracellular matrix proteins, respectively, and only one protein comes from Golgi apparatus membrane. The result indicated that bisecting GlcNAc modification preferred to occur in the proteins located at the surface of cells. In addition to tumor necrosis factor receptor superfamily member 11B [23], laminin subunit alpha-5 [36, 37], decorin [22], lysosome-associated membrane glycoprotein 2 [22], neprilysin [22], thy-1 membrane glycoprotein [23], nidogen-2 [22], integrin beta-1 [38, 39], cell surface glycoprotein MUC18 [23, 36], transferrin receptor protein 1 [39], integrin alpha-V [39], laminin subunit beta-2 [36, 37], tyrosine-protein kinase receptor UFO [36], immunoglobulin alpha-2 heavy chain [40, 41], carcinoembryonic antigen-related cell adhesion molecule 1 [36], fibronectin [36], other 25 glycoproteins are for the first time to be reported as ones possessing bisecting GlcNAc with confident MS data supports.

Figure 5 displays a typical MS/MS spectrum annotated as a glycopeptide KLHINHNNLTESVGPLPK, in which the N8 (asparagine) in NLT is glycosylated by a bisecting GlcNAc structure. The characteristic fragment ions that usually mark glycosylation presence, such as m/z 204.09 and m/z 366.14, are clearly observed in the spectrum due to their higher intensity. The evidences from its 5 peptide B ions (b3, b4, b5, b6 and b7) and 8 peptide Y ions (y2, y4, y5, y6, y7, y8, y9 and y10) support the peptide sequence identification and also the potential glycosylation modification location. The peaks at m/z 1391.71 (z=2) and m/z 1464.74 (z=2) matching to the [Pep+HexNAc_3_Hex] and [Pep+FucHexNAc_3_Hex] ions of the glycopeptide further prove that the glycosylation modification occurs on the N8 and contains a special bisecting GlcNAc structure, considering the ions specifically yielded by bisecting GlcNAc glycans. This glycopeptide is from lumican (accession number: P51884), and several studies have reported its vital roles on regulating tissue repair and embryonic development [42-45].

**Figure 5.**
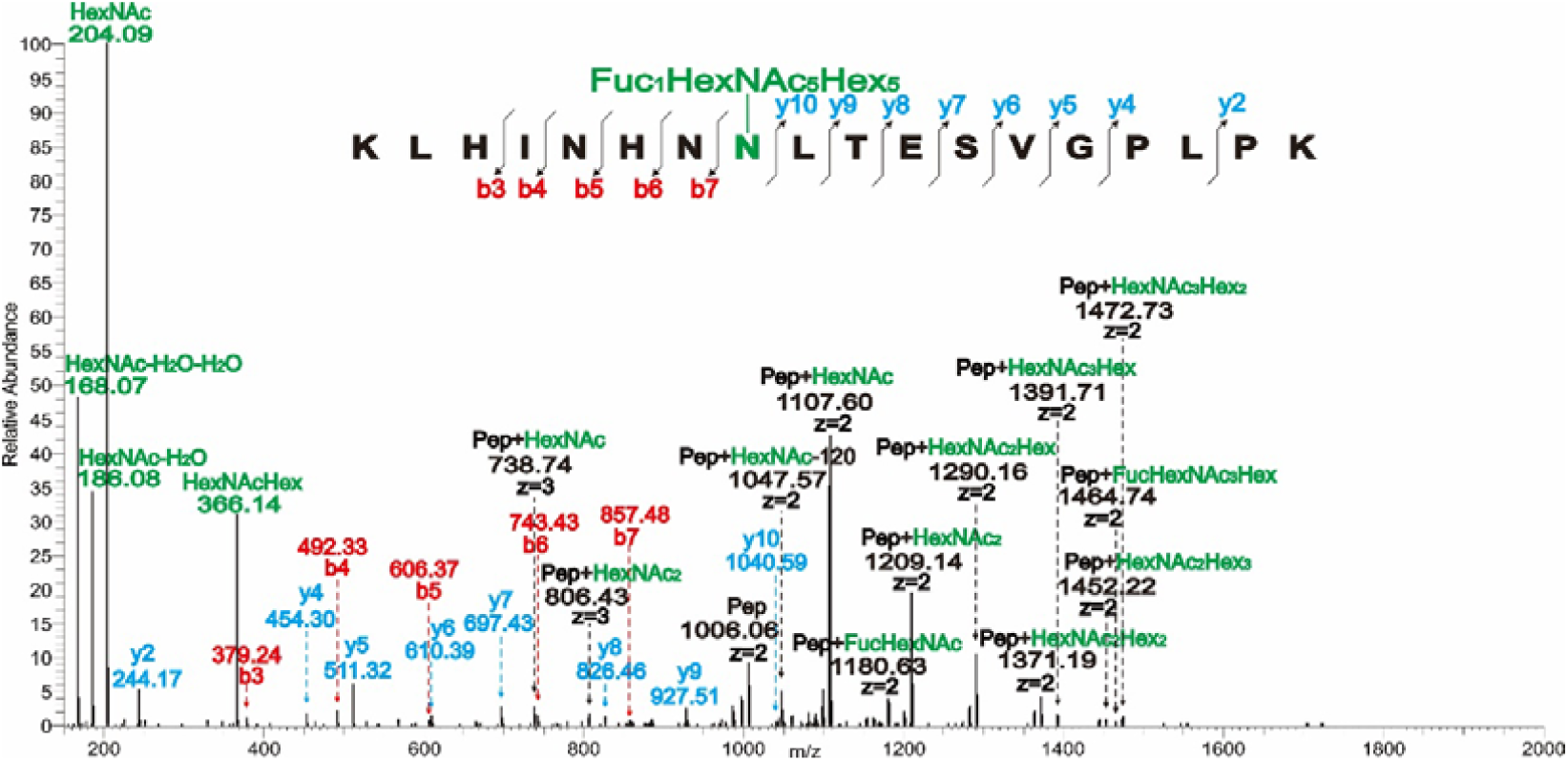
Annotated ESI MS/MS spectrum showing N-glycosylation on the N (asparagine) in the peptide KLHINHNNLTESVGPLPK. This Pep (peptide) is from lumican (P51884). All ions are [P+H]^+^ or [P+2H]^2+^ or [P+2H]^3+^. Double charged ions are annotated as z=2, triple charged ions are annotated as z=3, others are monocharged. The number indicated above the peak in the spectrum is the m/z value of the ion that has been detected by the mass spectrometer. To make the annotation clearer, ions are labelled in different colors.

The method that employed here in bisecting GlcNAc determination referenced a paper issued in *Analytical Chemistry* [22], which was designed to determine bisecting GlcNAc on glycopeptides by their characteristic ion(s) in fragmented MS/MS spectra under low energy collision. In this recently published paper, 25 glycoproteins (possessing bisecting GlcNAc) were identified from rat kidney tissue [22], 4 of which (Q01129 decorin, P17046 lysosome-associated membrane glycoprotein 2, P07861 neprilysin and B5DFC9 nidogen-2) were found to be protein analogues with those identified in our human amnion sample, more importantly, one of which, neprilysin, has the same bisecting GlcNAc location (site N285) as the human neprilysin (P08473) in our amnion sample.

Additionally, the same strategy was adopted to search the glycopeptides with bisecting GlcNAc structure from the proteomic data collected from human bladder, kidney and stomach, which had been released in our previous publication [46]. Without surprising, we confirmed that there was no bisecting GlcNAc structures in the corresponding peptides from human bladder and kidney proteins; and in human stomach there was only one glycopeptide HYTNSSQDVTVPC(Carbamidomethyl)R from human immunoglobulin alpha-2 heavy chain (P0DOX2), in which N-glycosylation occurred on the N in NSS, and its glycan is composed of FucHexNAc_5_Hex_5_. This result suggests some bisecting GlcNAc modifications might be specifically occurred in the proteins expressed in human amniotic membrane and essential in embryo development.

### Bioinformatic analyses

To investigate the functional roles of these glycoproteins with bisecting GlcNAc modification, gene ontology (GO) and Kyoto Encyclopedia of Genes and Genomes (KEGG) pathway analyses were perfomed. GO analysis results (Supplementary Figure 2) show that 7 of the glycoproteins (P02786, Q8NES3, P30530, P04216, P05556, Q07954 and O00300) are likely involved in immune system process. Pathway analysis results (Supplementary Figure 3) indicated that the pathways that the 41 glycoproteins are involved in can be classified into 5 categories: cellular processes, environmental information processing, human diseases, metabolism and organismal systems. In human disease category, 2 of the glycoproteins (P0DOX2 and P10321) are probably related to immune diseases, and in organismal system category, 8 of the glycoproteins (O75339, P02786, P04216, P05556, P08473, P0DOX2, P10321 and Q7Z7G0) may play important roles in immune system. Glycoprotein HLA class I histocompatibility antigen C alpha chain (P10321) is likely needs further investigation because it appears in both 2 categories.

## Conclusion

The method that employed in the glycomic analysis can determine the presence of bisecting GlcNAc on the proteins from a single tissue sample, and extra derivatization (e.g. partially methylated alditol acetates derivatization) or process (e.g. phaseolus vulgaris erythroagglutinin, β1,4-galactosyltransferase) is not required when using this method, it therefore will facilitate the studies of bisecting GlcNAc containing glycans greatly.

It should be noted that the two characteristic ions may not present in all MS2 spectra of bisecting glycopeptides, so it could miss some identification via this method. This method is recently published and now rapidly employed in our work, which makes this paper as a real-time update. Our glycomic and glycoproteomic analyses of human amniotic membrane for the first time showed that functional glycan structure that is present on human gametes and trophoblasts is also expressed on amniotic membrane, and this provides another evidence for the hu-FEDS hypothesis.

## Materials and methods

Chemical reagents were ordered from Sigma-Aldrich unless otherwise specified. Trypsin was bought from Promega. Peptide N-glycosidase F (PNGase F) was purchased from New England Biolabs. Reagents used for MSn (n=1-8), HPLC separation and LC-MS/MS were ordered from Thermo Fisher Scientific. The human placenta was kindly provided by Shenzhen Seventh People’s Hospital (Guangdong Province, China) with a signed informed consent form, which has been approved by the Ethics Committees in the hospital.

### Preparation of human amniotic membrane and determination of protein concentration

Human amniotic membrane was isolated from a placenta accordingly [30]. The protein extraction from amniotic membrane was performed as previously reported [31]. Protein concentration of homogenized amniotic membrane was determined by Bradford assay and SDS-PAGE was performed to verify the rationality of the protein concentration. Proteins from the membrane were extracted following our lab protocol.

### Processing of human amniotic membrane to acquire N-glycans

The membrane was subjected to a standard protocol [26]. Briefly, the membrane was suspended in lysis buffer before homogenization and sonication were performed. The homogenates were reduced and carboxymethylated and then dialyzed against a 50 mM ammonia bicarbonate buffer, pH 7.5, after which the sample was lyophilized, and then treated with trypsin. The treated sample was purified using a C18 Waters cartridge prior to the release of N-glycans by PNGase F digestion. Released N-glycans were permethylated and then purified using a Sep-Pak C18 cartridge (Waters) prior to MS analysis.

### MSn (n=1-8) Analysis for glycomics

MS data were obtained by using a Bruker Ultraflextreme MALDI-TOF/TOF mass spectrometer. Purified permethylated glycans were dissolved in 20 μL methanol, and 1 μL of the sample was mixed with 1 mL of matrix, 20 mg/mL 2,5-dihydroxybenzoic acid (DHB) in 70% (v/v) aqueous methanol and loaded on to a metal target plate. The instrument was run in the reflectron positive ion mode. MSn (n=2-8) data were acquired using a Thermo Scientific Orbitrap Fusion Lumos Tribrid mass spectrometer via direct infusion of the sample dissolving in 100 μL of 1 mM NaOH in 50% methanol (infusion buffer) into the mass spectrometer.

### HPLC separation for glycoproteomics

The peptides were desalted and then dissolved in the high concentration organic solutions with shaking for 15 minutes at room temperature and then separated by centrifuge for 10 minutes at 16,000g. For further HPLC separation details please see our previously published paper [31].

### LC-MS/MS

The peptides were passed onto a Thermo Scientific Orbitrap Fusion Lumos Tribrid Mass Spectrometer for protein identification coupling with a RP C18 column with a LC gradient (5-25% buffer B for 95minutes, 25-30% for 10minutes, 30-80% for 5minutes). The MS parameters were set as before with a 30% NCE. Considering an additionally dimensional separation of the peptides into supernatant and pellets based on their hydrophobicity, we modified the LC gradient for the first 4 and last 4 peptide fractions from high pH RP column for more identification. For fraction 1-4, more shallow gradient at low concentration of ACN were adopted to improve the hydrophilic peptide identification (5-18% buffer B for 95minutes, 18-35% for 10minutes, 35-80% for 5minutes) and oppositely the gradient (10-26% buffer B for 95minutes, 26-35% for 10minutes, 35-80% for 5minutes) suit for the more hydrophobic peptide identification were used for the fractions of 17-20 to get more hydrophobic peptide identification. Each fraction was injected twice for more confident identification.

### Database analysis

The glycomic data were analyzed using Xcalibur and GlycoWorkbench to get glycan structure. To verify the structure, the MSn (n=2-8) spectra related to bisecting GlcNAc containing glycan were manually checked. The proteomic data was searched by Proteome discoverer against the Swiss-prot human database. The false discovery rate was less than 1% at both PSM and protein level during searching and automatically calculated by the software [47, 48]. The glycoproteomic data was processed by the combination of Proteome Discoverer and Xcalibur to get the glycosylation site and the corresponding glycan structure. To confirm the identification, the MS/MS spectra of glycans and glycopeptides were manually checked for oxonium ions, monosaccharide and oligosaccharide neutral loss patterns.

### Bioinformatic analysis for the glycoproteins containing bisecting GlcNAc

The Gene Ontology (GO) annotation was performed using NCBI non-redundant (nr) database and software Blast2GO. Proteins were categorized into molecular function, cellular component and biological process according to Gene Ontology (GO) terms. Kyoto Encyclopedia of Genes and Genomes (KEGG, version 89.1) as well as software Blast2KO were utilized to annotate pathways. The P<0.05 was considered significant (Kall et al., 2007; Perkins et al., 1999; Tatusov et al., 2000).

## Authors’ contributions

QC participated in project design, carried out glycomic studies, participated in glycoproteomic data analysis and drafted the manuscript. YZ carried out glycoproteomic studies and data analysis. KZ participated in glycomic and glycoproteomic data analysis. JL participated in glycoproteomic data analysis. HP participated in glycoproteomic data analysis. XW participated in glycoproteomic data analysis. SL participated in glycoproteomic studies. DH participated in glycomic studies. ZL participated in glycoproteomic studies. YZ participated in glycoproteomic studies. GH participated in glycomic studies. FG participated in glycomic studies and discussion. HL provided the sample and participated in project discussion. SL participated in project design. YR participated in project design and drafted the manuscript. All authors read and approved the final manuscript.

## Acknowledgements

This work was supported by the National Natural Science Foundation of China (Number:31500670 and Number: 31700728).

## Supplementary data

**Supplementary Figure 1.**
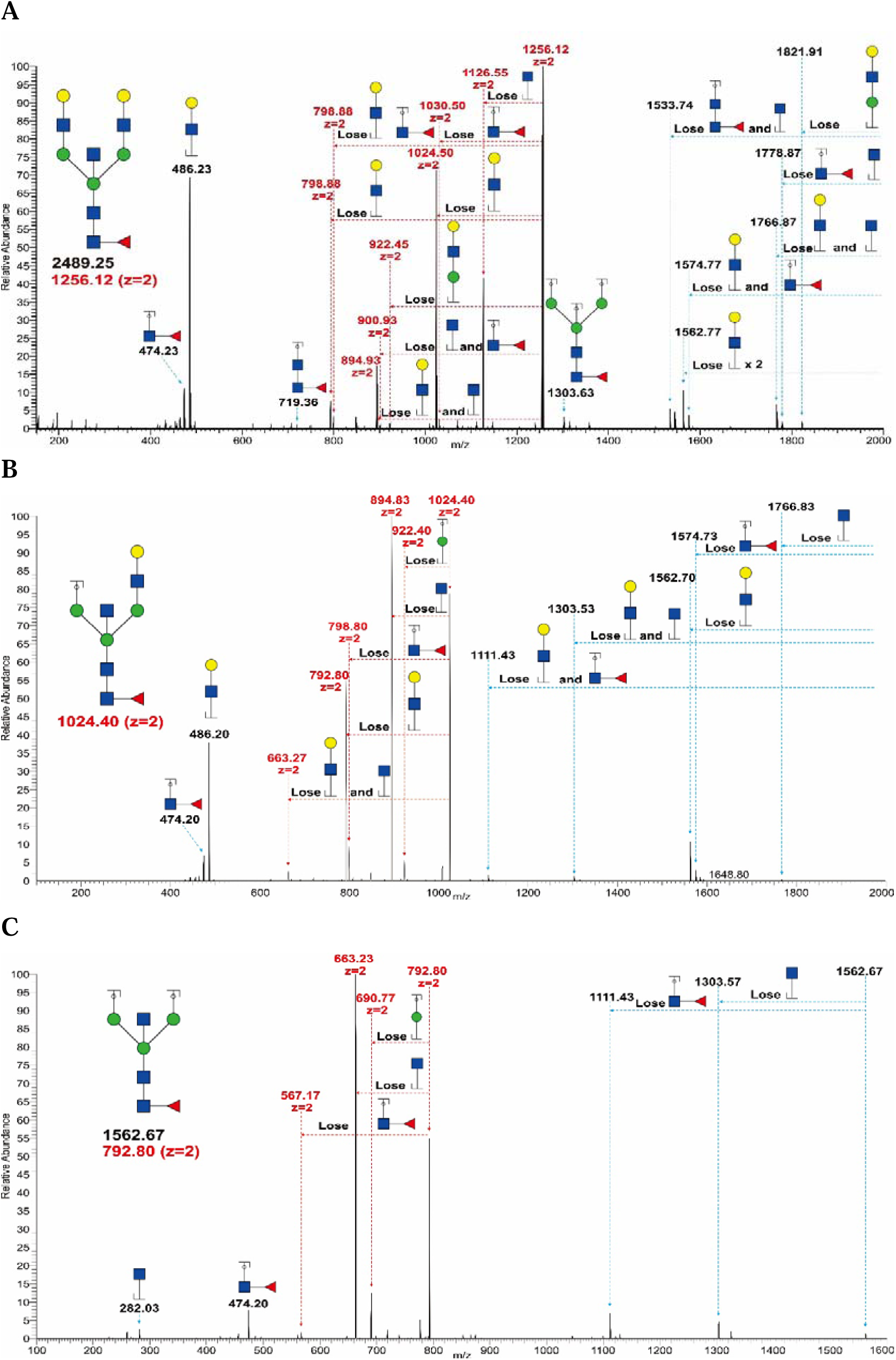
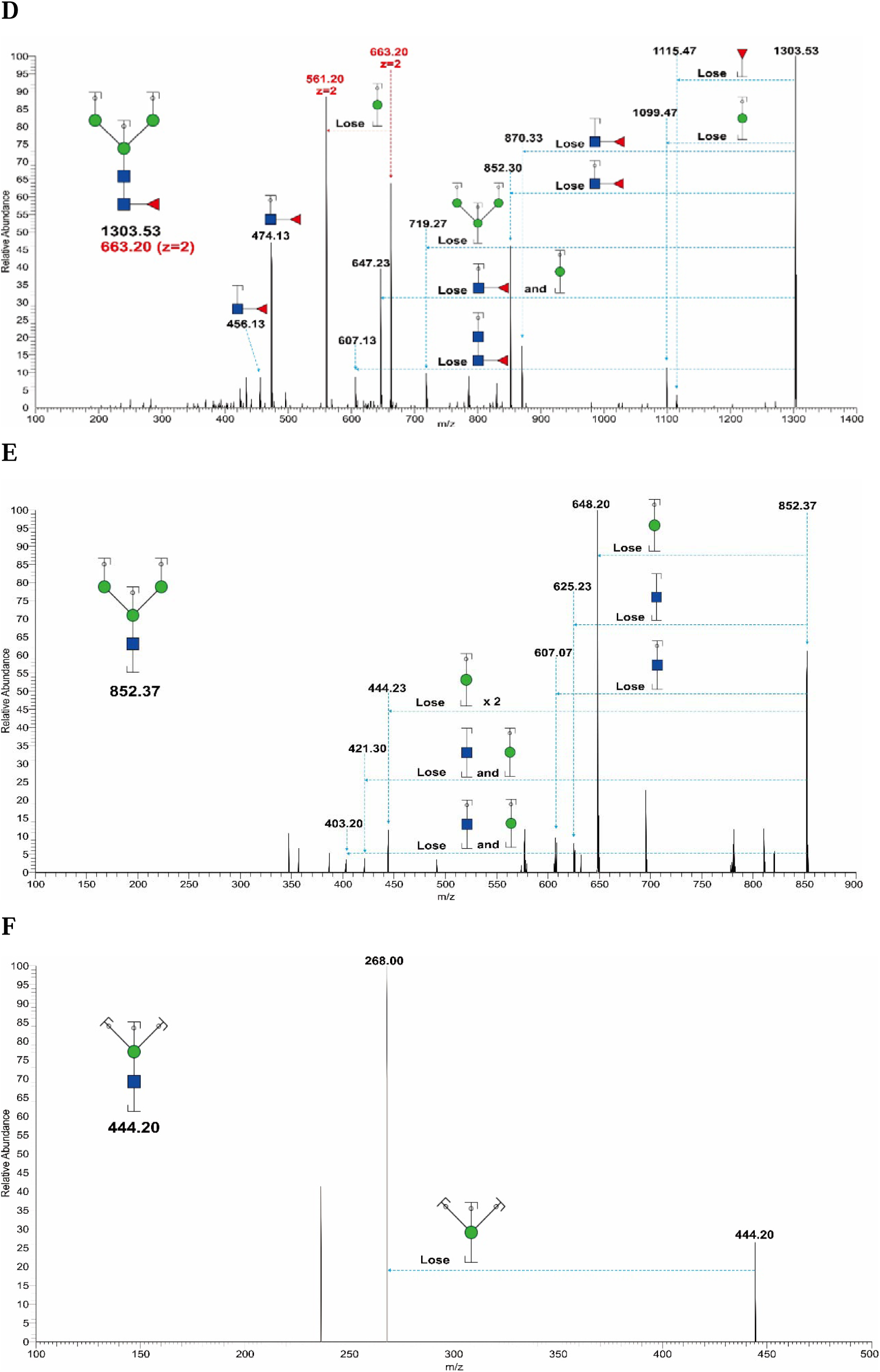
A. annotated ESI FTMS MS2 spectrum of permethylated N-glycan at m/z 2489.25; B. annotated ESI ITMS MS3 spectrum of bicharged permethylated N-glycan fragment ion at m/z 1024.4; C. annotated ESI ITMS MS4 spectrum of bicharged permethylated N-glycan fragment ion at m/z 792.8; D. annotated ESI ITMS MS5 spectrum of monocharged permethylated N-glycan fragment ion at m/z 1303.53; E. annotated ESI ITMS MS6 spectrum of monocharged permethylated N-glycan fragment ion at m/z 852.37; F. annotated ESI ITMS MS8 spectrum of monocharged permethylated N-glycan fragment ion at m/z 444.2 from human amniotic membrane. Assignments of the possible fragment ions are indicated on the cartoons and on the spectrum. Ions with different charges are labeled in different colors. The number indicated above the peak is the m/z value of the fragment ion (resulting ion) that has been detected by the mass spectrometer. Data were acquired in the form of [M+Na]^+^ or [M+2Na]^2+^. ▪ GlcNAc, • Man, • Gal, ▴ Fuc.

**Supplementary Figure 2.**
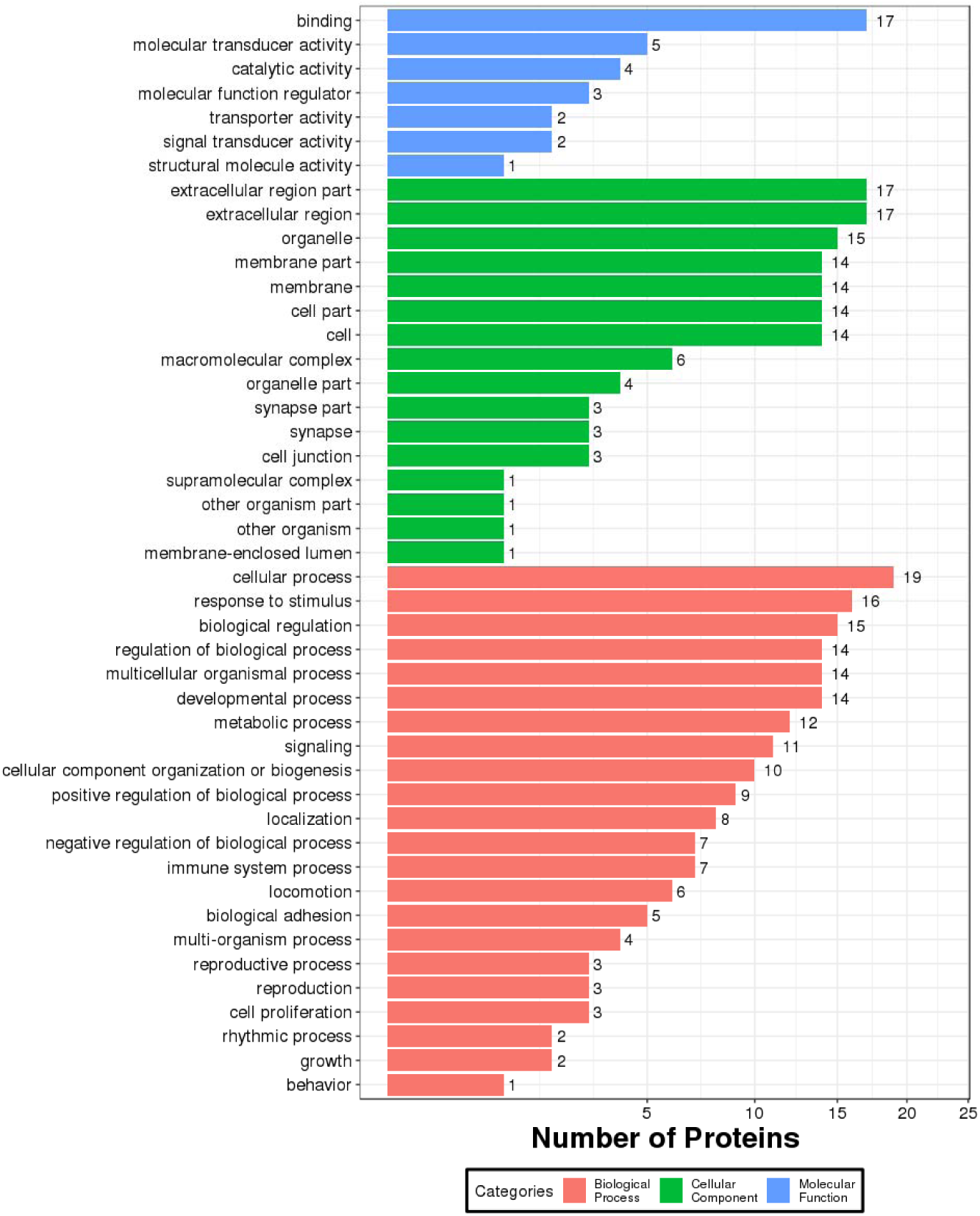
The distribution of identified bisecting GlcNAc containing glycoproteins in molecular function, cellular component and biological process. One protein may be involved in multiple categories

**Supplementary Figure 3.**
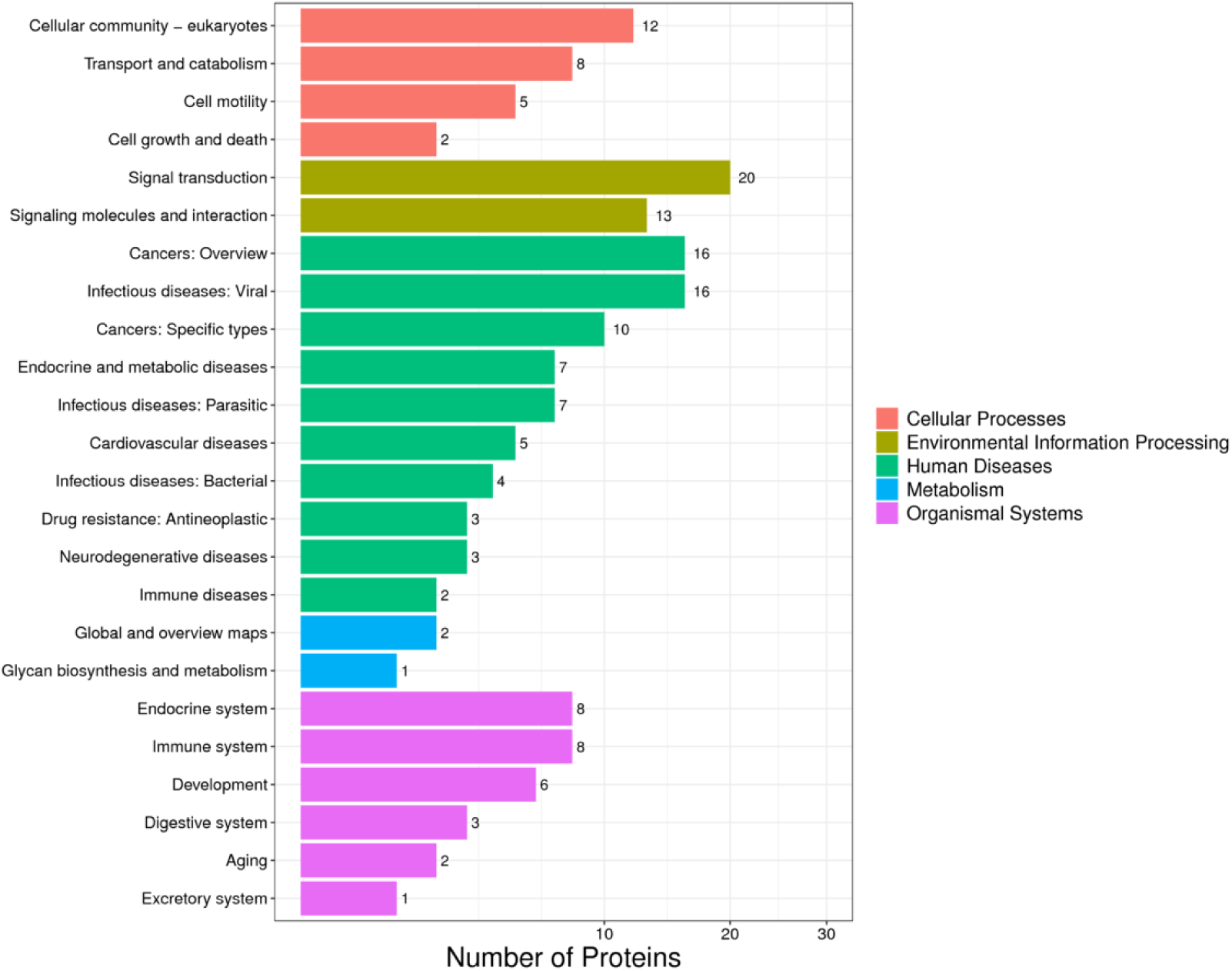
KEGG analyses result of the identified bisecting GlcNAc containing glycoproteins.

**Supplementary Table 1.**
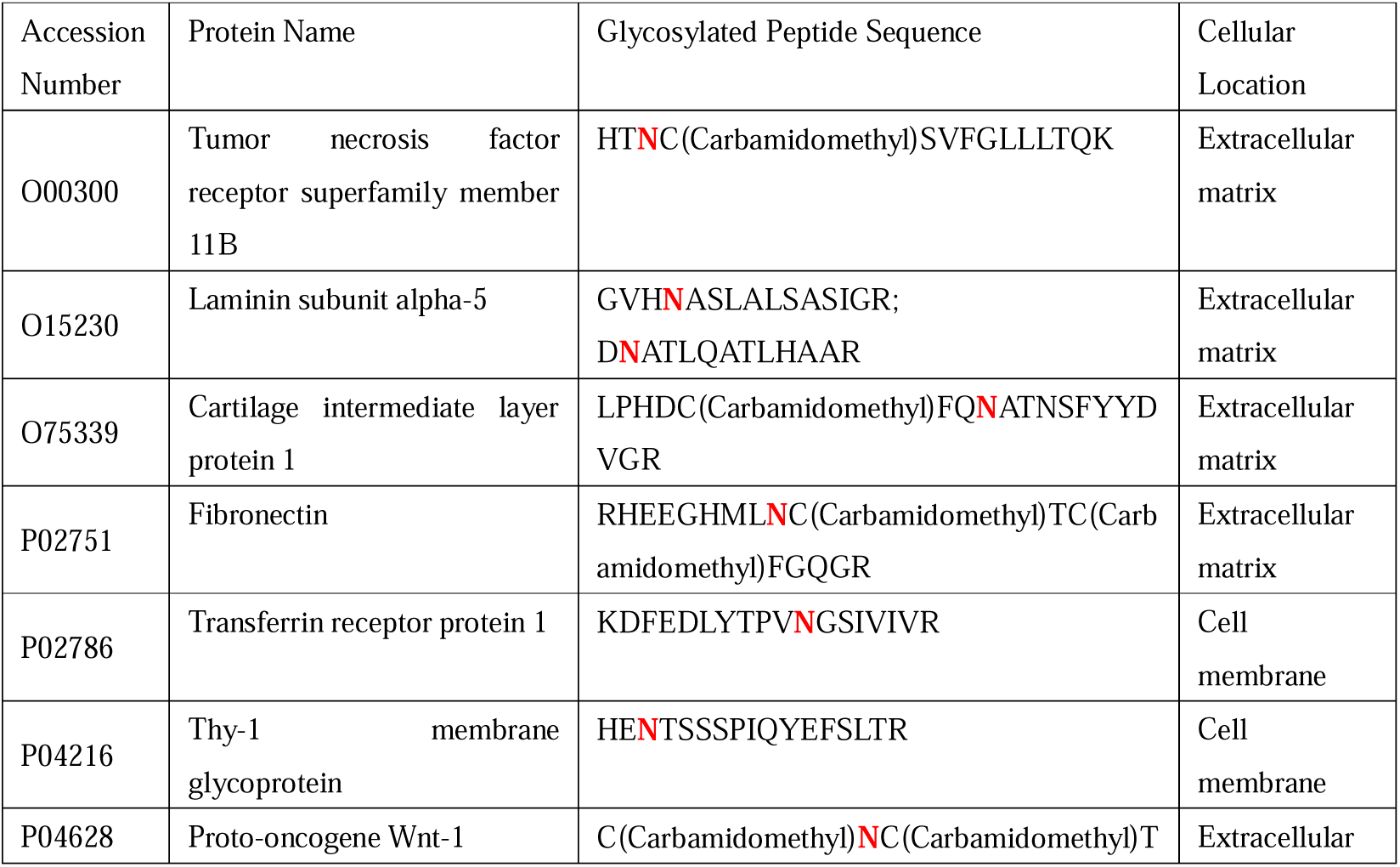
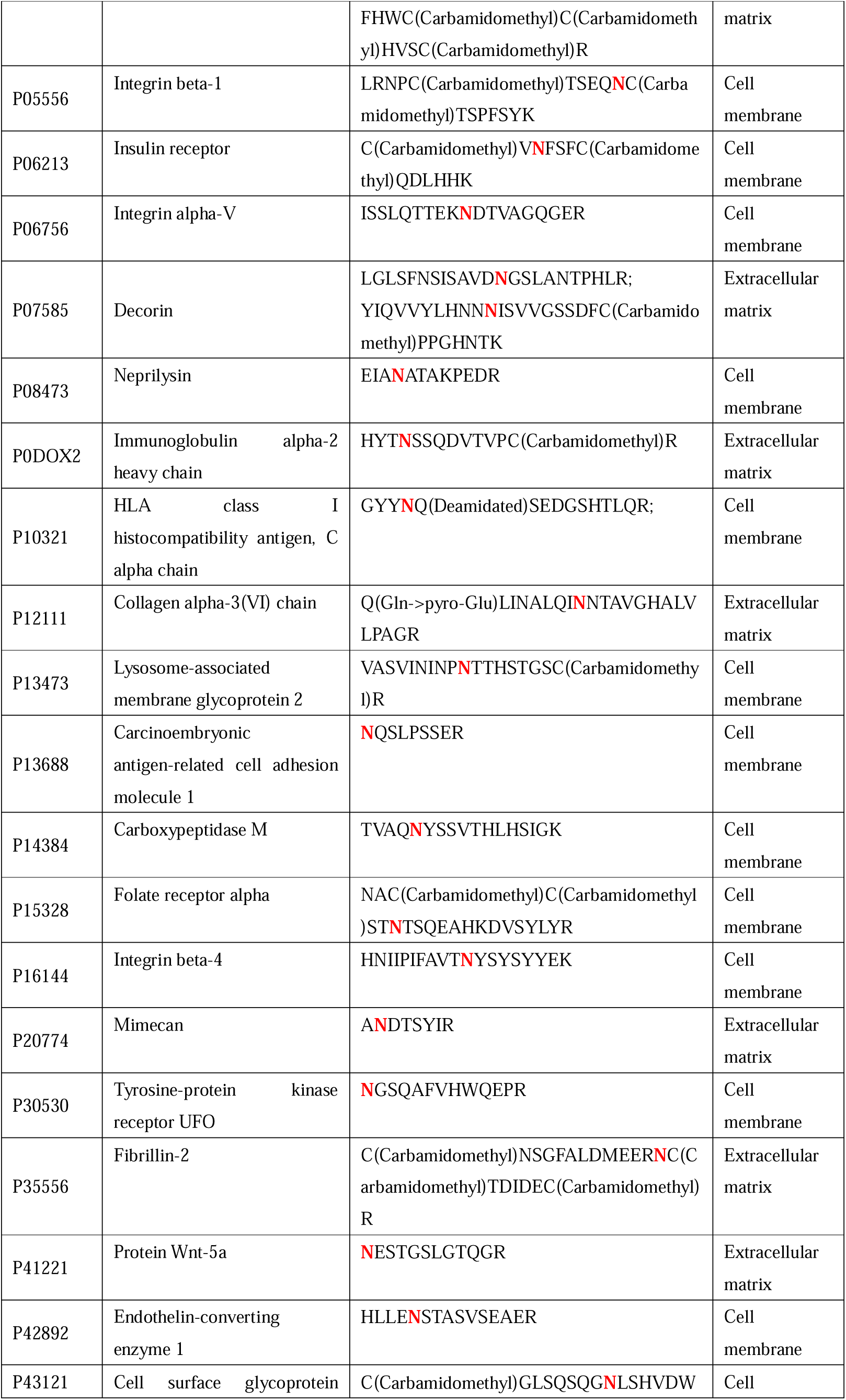
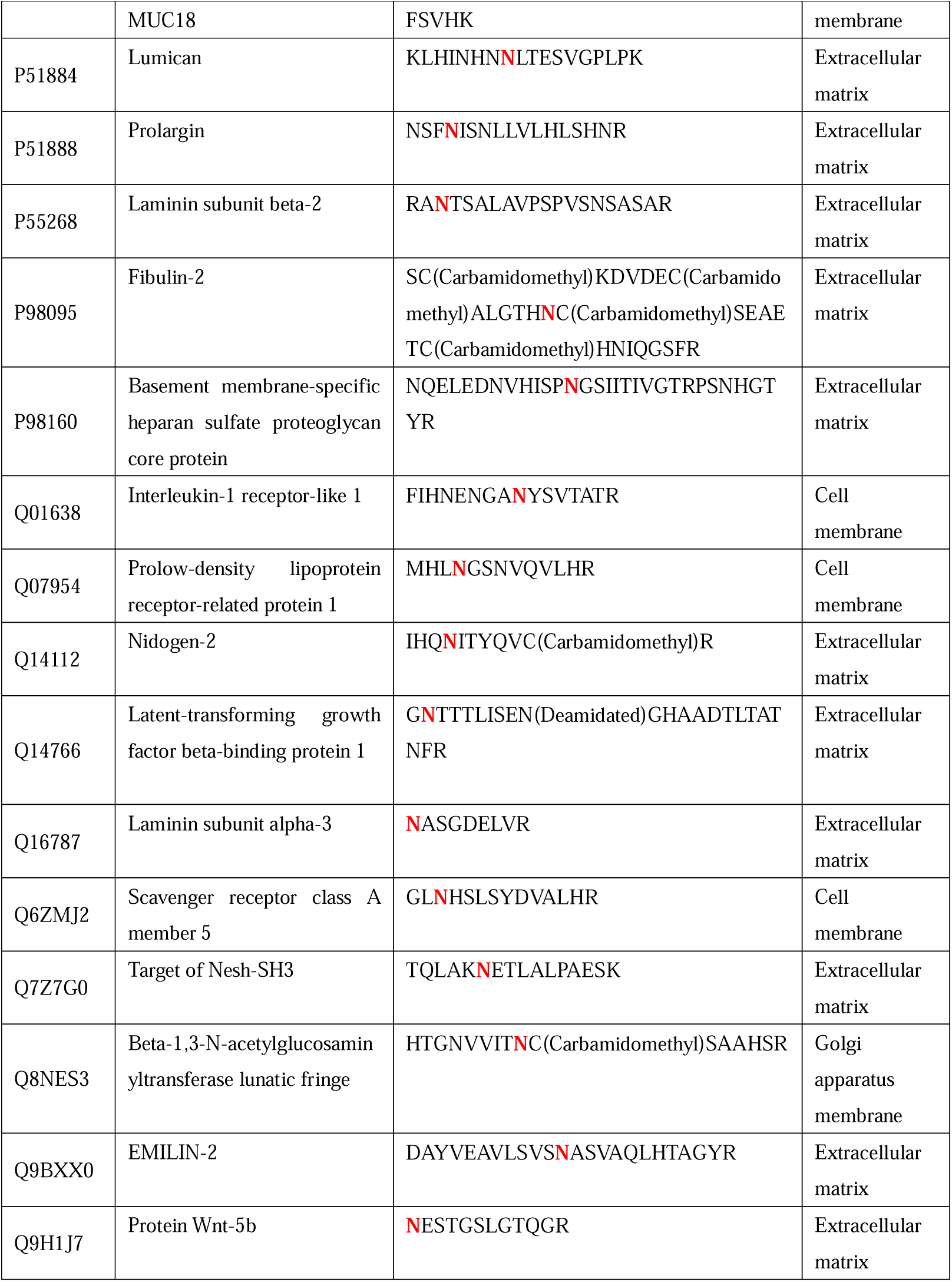
Accession numbers, protein names, one glycosylated peptide sequence of all identified glycoproteins possessing bisecting GlcNAc and cellular location of each glycoprotein. N-glycosylation site was emphasized using red color, cellular location was obtained via searches in UniProt library

## References

1. Schachter H. The ‘yellow brick road’ to branched complex N-glycans. Glycobiology. 1991;1(5):453–61.

2. Kizuka Y, Taniguchi N. Neural functions of bisecting GlcNAc. Glycoconj J. 2018;35(4):345–51.

3. Varki A. Essentials of glycobiology. 2nd ed. Cold Spring Harbor, N.Y.: Cold Spring Harbor Laboratory Press; 2009. xxix, 784 p. p.

4. Harpaz N, Schachter H. Control of glycoprotein synthesis. Processing of asparagine-linked oligosaccharides by one or more rat liver Golgi alpha-D-mannosidases dependent on the prior action of UDP-N-acetylglucosamine: alpha-D-mannoside beta 2-N-acetylglucosaminyltransferase I. J Biol Chem. 1980;255(10):4894–902.

5. Yamashita K, Tachibana Y, Kobata A. The structures of the galactose-containing sugar chains of ovalbumin. J Biol Chem. 1978;253(11):3862–9.

6. Chen Q, Pang PC, Cohen ME, Longtine MS, Schust DJ, Haslam SM, et al. Evidence for Differential Glycosylation of Trophoblast Cell Types. Mol Cell Proteomics. 2016;15(6):1857–66.

7. Miwa HE, Song Y, Alvarez R, Cummings RD, Stanley P. The bisecting GlcNAc in cell growth control and tumor progression. Glycoconj J. 2012;29(8-9):609–18.

8. Narasimhan S. Control of glycoprotein synthesis. UDP-GlcNAc:glycopeptide beta 4-N-acetylglucosaminyltransferase III, an enzyme in hen oviduct which adds GlcNAc in beta 1-4 linkage to the beta-linked mannose of the trimannosyl core of N-glycosyl oligosaccharides. J Biol Chem. 1982;257(17):10235–42.

9. Schachter H. Complex N-glycans: the story of the “yellow brick road“. Glycoconj J. 2014;31(1):1–5.

10. Nagae M, Yamanaka K, Hanashima S, Ikeda A, Morita-Matsumoto K, Satoh T, et al. Recognition of bisecting N-acetylglucosamine: structural basis for asymmetric interaction with the mouse lectin dendritic cell inhibitory receptor 2. J Biol Chem. 2013;288(47):33598–610.

11. Yoshimura M, Nishikawa A, Ihara Y, Taniguchi S, Taniguchi N. Suppression of lung metastasis of B16 mouse melanoma by N-acetylglucosaminyltransferase III gene transfection. Proc Natl Acad Sci U S A. 1995;92(19):8754–8.

12. Yang W, Ramadan S, Orwenyo J, Kakeshpour T, Diaz T, Eken Y, et al. Chemoenzymatic synthesis of glycopeptides bearing rare N-glycan sequences with or without bisecting GlcNAc. Chem Sci. 2018;9(43):8194–206.

13. Patankar MS, Ozgur K, Oehninger S, Dell A, Morris H, Seppala M, et al. Expression of glycans linked to natural killer cell inhibition on the human zona pellucida. Mol Hum Reprod. 1997;3(6):501–5.

14. Yoshimura M, Ihara Y, Ohnishi A, Ijuhin N, Nishiura T, Kanakura Y, et al. Bisecting N-acetylglucosamine on K562 cells suppresses natural killer cytotoxicity and promotes spleen colonization. Cancer Res. 1996;56(2):412–8.

15. el Ouagari K, Teissie J, Benoist H. Glycophorin A protects K562 cells from natural killer cell attack. Role of oligosaccharides. J Biol Chem. 1995;270(45):26970–5.

16. Clark GF, Oehninger S, Patankar MS, Koistinen R, Dell A, Morris HR, et al. A role for glycoconjugates in human development: the human feto-embryonic defence system hypothesis. Hum Reprod. 1996;11(3):467–73.

17. Pang PC, Haslam SM, Dell A, Clark GF. The human fetoembryonic defense system hypothesis: Twenty years on. Mol Aspects Med. 2016;51:71–88.

18. Kui Wong N, Easton RL, Panico M, Sutton-Smith M, Morrison JC, Lattanzio FA, et al. Characterization of the oligosaccharides associated with the human ovarian tumor marker CA125. J Biol Chem. 2003;278(31):28619–34.

19. Pang PC, Tissot B, Drobnis EZ, Sutovsky P, Morris HR, Clark GF, et al. Expression of bisecting type and Lewisx/Lewisy terminated N-glycans on human sperm. J Biol Chem. 2007;282(50):36593–602.

20. Chen Q. Mass spectrometric investigation of biomedically important glycosylation: Imperial College London; 2015.

21. Monnich M, Eller S, Karagiannis T, Perkams L, Luber T, Ott D, et al. Highly Efficient Synthesis of Multiantennary Bisected N-glycans Based on Imidates. Angew Chem Int Ed Engl. 2016;55(35):10487–92.

22. Dang L, Shen J, Zhao T, Zhao F, Jia L, Zhu B, et al. Recognition of Bisecting N-Glycans on Intact Glycopeptides by Two Characteristic Ions in Tandem Mass Spectra. Anal Chem. 2019;91(9):5478–82.

23. Allam H, Aoki K, Benigno BB, McDonald JF, Mackintosh SG, Tiemeyer M, et al. Glycomic analysis of membrane glycoproteins with bisecting glycosylation from ovarian cancer tissues reveals novel structures and functions. J Proteome Res. 2015;14(1):434–46.

24. Pang PC, Chiu PC, Lee CL, Chang LY, Panico M, Morris HR, et al. Human sperm binding is mediated by the sialyl-Lewis(x) oligosaccharide on the zona pellucida. Science. 2011;333(6050):1761–4.

25. Dell A, Morris HR. Glycoprotein structure determination by mass spectrometry. Science. 2001;291(5512):2351–6.

26. North SJ, Jang-Lee J, Harrison R, Canis K, Ismail MN, Trollope A, et al. Mass spectrometric analysis of mutant mice. Methods Enzymol. 2010;478:27–77.

27. Gupta A, Kedige SD, Jain K. Amnion and Chorion Membranes: Potential Stem Cell Reservoir with Wide Applications in Periodontics. Int J Biomater. 2015;2015:274082.

28. Toda A, Okabe M, Yoshida T, Nikaido T. The potential of amniotic membrane/amnion-derived cells for regeneration of various tissues. J Pharmacol Sci. 2007;105(3):215–28.

29. Rock T, Bosmuller HC, Bartz-Schmidt KU, Rock D. Surgical management of a conjunctival nevus with amniotic membrane transplantation. Int Med Case Rep J. 2018;11:161–5.

30. Sundell IB, Koka P. Placental membrane as a source of mesenchymal stem cells. J Stem Cells. 2010;5(2):83–8.

31. Lin Z, Zhang Y, Pan H, Hao P, Li S, He Y, et al. Alternative Strategy To Explore Missing Proteins with Low Molecular Weight. J Proteome Res. 2019;18(12):4180–8.

32. Caldwell RL, Caprioli RM. Tissue profiling by mass spectrometry: a review of methodology and applications. Mol Cell Proteomics. 2005;4(4):394–401.

33. Antonopoulos A, Demotte N, Stroobant V, Haslam SM, van der Bruggen P, Dell A. Loss of effector function of human cytolytic T lymphocytes is accompanied by major alterations in N- and O-glycosylation. J Biol Chem. 2012;287(14):11240–51.

34. Savaryn JP, Toby TK, Kelleher NL. A researcher’s guide to mass spectrometry-based proteomics. Proteomics. 2016;16(18):2435–43.

35. McAlister GC, Phanstiel D, Wenger CD, Lee MV, Coon JJ. Analysis of tandem mass spectra by FTMS for improved large-scale proteomics with superior protein quantification. Anal Chem. 2010;82(1):316–22.

36. Liu TH, Liu DH, Mo CJ, Sun L, Liu XX, Li W, et al. Glycosylation patterns and PHA-E-associated glycoprotein profiling associated with early hepatic encephalopathy in Chinese hepatocellular carcinoma patients. Am J Transl Res. 2016;8(10):4250–64.

37. Kariya Y, Kato R, Itoh S, Fukuda T, Shibukawa Y, Sanzen N, et al. N-Glycosylation of laminin-332 regulates its biological functions. A novel function of the bisecting GlcNAc. J Biol Chem. 2008;283(48):33036–45.

38. Zhang M, Wang M, Gao R, Liu X, Chen X, Geng Y, et al. Altered beta1,6-GlcNAc and bisecting GlcNAc-branched N-glycan on integrin beta1 are associated with early spontaneous miscarriage in humans. Hum Reprod. 2015;30(9):2064–75.

39. Park DD, Xu G, Wong M, Phoomak C, Liu M, Haigh NE, et al. Membrane glycomics reveal heterogeneity and quantitative distribution of cell surface sialylation. Chem Sci. 2018;9(29):6271–85.

40. Bartsch YC, Rahmoller J, Mertes MMM, Eiglmeier S, Lorenz FKM, Stoehr AD, et al. Sialylated Autoantigen-Reactive IgG Antibodies Attenuate Disease Development in Autoimmune Mouse Models of Lupus Nephritis and Rheumatoid Arthritis. Front Immunol. 2018;9:1183.

41. Epp A, Sullivan KC, Herr AB, Strait RT. Immunoglobulin Glycosylation Effects in Allergy and Immunity. Curr Allergy Asthma Rep. 2016;16(11):79.

42. Yeh LK, Chen WL, Li W, Espana EM, Ouyang J, Kawakita T, et al. Soluble lumican glycoprotein purified from human amniotic membrane promotes corneal epithelial wound healing. Invest Ophthalmol Vis Sci. 2005;46(2):479–86.

43. Coulson-Thomas VJ, Coulson-Thomas YM, Gesteira TF, Andrade de Paula CA, Carneiro CR, Ortiz V, et al. Lumican expression, localization and antitumor activity in prostate cancer. Exp Cell Res. 2013;319(7):967–81.

44. Cornuet PK, Blochberger TC, Hassell JR. Molecular polymorphism of lumican during corneal development. Invest Ophthalmol Vis Sci. 1994;35(3):870–7.

45. Grover J, Chen XN, Korenberg JR, Roughley PJ. The human lumican gene. Organization, chromosomal location, and expression in articular cartilage. J Biol Chem. 1995;270(37):21942–9.

46. Li S, He Y, Lin Z, Xu S, Zhou R, Liang F, et al. Digging More Missing Proteins Using an Enrichment Approach with ProteoMiner. J Proteome Res. 2017;16(12):4330–9.

47. Brosch M, Yu L, Hubbard T, Choudhary J. Accurate and sensitive peptide identification with Mascot Percolator. J Proteome Res. 2009;8(6):3176–81.

48. Lee SE, Song J, Bosl K, Muller AC, Vitko D, Bennett KL, et al. Proteogenomic Analysis to Identify Missing Proteins from Haploid Cell Lines. Proteomics. 2018;18(8):e1700386.

